# Could a female athlete run a 4-minute mile with improved aerodynamic drafting?

**DOI:** 10.1101/2024.08.15.608172

**Authors:** Edson Soares da Silva, Wouter Hoogkamer, Shalaya Kipp, Rodger Kram

**Author notes:** Corresponding Author: Rodger Kram, Integrative Locomotion Lab, University of Colorado, Boulder, United States of America.

## Abstract

In 2023, Faith Kipyegon set the female world record for running one mile (4:07.64). Here, we quantitively explore if improved aerodynamic drafting could allow her to run just 3.19% faster and thus break the 4-minute mile barrier. Drafting involves other athletes (pacers) running in formation around a designated athlete to reduce the aerodynamic drag force acting on the designated runner. Drafting allows the designated athlete to run faster at the same rate of metabolic energy consumption. Our overall approach was to estimate Kipyegon’s metabolic energy consumption during her mile world record performance. Then, we used empirically established relationships between horizontal resistive force, running velocity and metabolic power to estimate how much faster she could run at the same metabolic power if the aerodynamic force was reduced via drafting. Our calculations suggest that Kipyegon could run ∼3:59.37 with drafting provided by one pacer in front and one in back who change out with two other pacers at 800 m.

## 1. Introduction

In 1954, two milestones in track and field history were reached. On May 6, Roger Bannister became the first human to run one mile (1609.344 m) in under 4 minutes (3:59.4) (Bannister, 1955). Shortly thereafter, Diane Leather ran a mile in 4:59.6, becoming the first female athlete to run under 5 minutes (Fountain, 2024). Over the ensuing seven decades, the female mile record continued to progress and in 2023, Faith Kipyegon, ran 4:07.64 (World Athletics, 2023). In this paper, we quantitively explore if improved aerodynamic drafting could allow a top female athlete, like Kipyegon, to run just 3.19% faster and thus break the 4-minute mile barrier.

Aerodynamic drafting occurs when other athletes (pacers) run in formation around a designated athlete to reduce the aerodynamic drag force acting on the designated runner (Hoogkamer et al., 2017). Drafting allows the designated athlete to run at a given speed with a slower rate of metabolic energy consumption. Or, more relevant here, drafting allows the designated athlete to run faster at the same rate of metabolic energy consumption (Kipp et al., 2019). When Bannister first broke the 4-minute mile, he drafted closely behind two different pacers for more than 80% of the race. In contrast, during her world record mile race, Kipyegon ran behind pacers for just the first ∼900 m (∼56% of the race) and thereafter ran solo, with no drafting. Further, she did not consistently run closely behind her pacers and thus the drafting was suboptimal.

Scientists and engineers have quantified the air resistance force experienced by a solo runner in several ways: scaled manikins in wind tunnels (Hill, 1928; Marro et al., 2023), actual human runners on treadmills in a wind tunnel (Pugh, 1971; Davies, 1980; Pecchiari et al., 2023) and using computational fluid dynamics simulations (CFD) (Beves & Ferguson, 2017; Blocken, 2019; Polidori et al., 2020; Schickhofer & Hanson, 2021). The classic equation (Rayleigh, 1876) for calculating aerodynamic drag force (F_aero_) is:

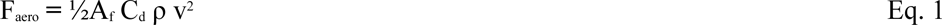

where F_aero_ is in units of newtons, A_f_ is the frontal area of the runner in m^2^, C_d_ is the dimensionless coefficient of drag, ρ is air density in kg/m^3^ and v is running velocity in m/s. Using Equation 1, Bannister’s height (1.88 m) and body mass (70 kg) along with anthropometric equations*, and reasonable values for C_d_ (0.9) and air density 1.2 kg/m^3^, we calculate that at 4-minute mile pace (6.706 m/s), the aerodynamic drag force acting on him when running solo was approximately 12.58 N or 1.83% of his body weight (BW). For the smaller Kipyegon (1.57 m and 42 kg), the corresponding values are 8.89 N and 2.16% of BW, notably a greater percentage of her body weight due to her greater surface area:volume ratio.

Classic (Pugh, 1971) and recent (Valsecchi et al., 2022; da Silva et al., 2022) empirical physiological measurements concur that the rate of metabolic energy consumption (i.e., metabolic power) required to overcome horizontal resistive forces comprises approximately 6% of the total metabolic power required to run per BW of resistive force.

Using da Silva et al.’s average value of 6.13% per BW, and the values in the previous paragraph, we calculate that overcoming air resistance when running solo at 4-minute mile pace comprised 11.4% of Bannister’s metabolic power and the corresponding value for Kipyegon is 13.5%. Wind tunnel and CFD methods can estimate how much various drafting formations of pacers reduce the drag force. Estimates of drag force reduction (drafting effectiveness) range from 9.7 to 85% depending on the pacer formation, spacing and measurement/simulation methods (Pugh, 1970, 1971; Beves & Ferguson, 2017; Schickhofer & Hanson, 2021; Beaumont et al., 2021; Polidori et al., 2020; Blocken, 2019; Valsecchi et al., 2022). Since middle-distance running performance is largely determined by an athlete’s ability to supply metabolic energy to their muscles (Brandon, 1995), reducing aerodynamic drag and hence the rate of metabolic energy required, can enhance performance considerably (Kipp et al., 2019).

We hypothesized that given sufficient aerodynamic drafting, an elite female athlete could run a 4-minute mile. Our specific goal in this paper was to model and calculate what drafting effectiveness would be needed.

## 2. Methods

Our overall approach was to first estimate Kipyegon’s the total rate of metabolic energy consumption (i.e. metabolic power) during each lap of her mile world-record performance. Then, we used our empirically established relationship between horizontal resistive force and metabolic power (da Silva et al., 2022) to estimate how much faster she could run at the same metabolic power if the aerodynamic force was reduced via drafting. Middle-distance running speeds require the supply of metabolic energy to the muscles via both oxidative (aerobic) and non-oxidative (glycolytic) metabolic pathways (Spencer & Gastin, 2001; Duffield et al., 2005). Using indirect respirometry, it is possible to accurately measure the oxidative metabolic rate at sustainable running speeds. But, measuring the rate of non-oxidative metabolism during high-speed running would require radioactive tracers (Brooks et al. 2005) and to our knowledge, has not yet been done. One can, however, extrapolate from the rates of oxidative metabolism measured at slower speeds to estimate the total rate of metabolic energy consumption (metabolic power) at faster speeds.

Batliner et al., (2018) measured the oxidative metabolic power (P_met_) in watts per kilogram body mass required for treadmill (tm) running in sub-elite athletes over a range of running speeds (1.78 – 5.14 m/s). Their curvilinear regression equation is:

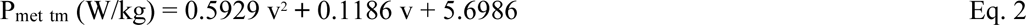

where v is running velocity in units of m/s. Note that during treadmill running there is no significant aerodynamic force to overcome.

We then used da Silva et al.’s (2022) finding that for each 1% of BW of horizontal impeding force, metabolic power during running is, on average, 6.13% greater than during normal treadmill running. Conveniently, the da Silva et al. value applies at different running speeds. Thus, for solo overground (og) running, total P_met_ (in W/kg) is equal to:

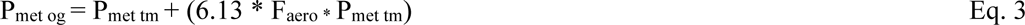

where F_aero_ is expressed as decimal BW, i.e. 1% of BW is 0.01. Equation 3 can be modified to account for the drafting effectiveness (eff_draft_) where 0.50 would represent a 50% reduction in drag force.

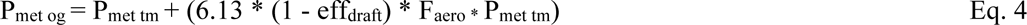

Substituting Equations 2 and 3 into Equation 4 yields:

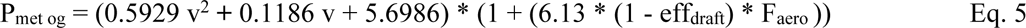

By calculating F_aero_ for a solo runner and assuming a certain eff_draft_, Equation 5 reduces to a quadratic equation in terms of v:

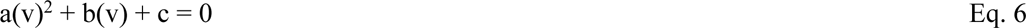

where a is equal to: 0.5929 * (1 + (6.13 * (1 - eff_draft_) * F_aero_))

b is equal to: 0.1186 * (1 + (6.13 * (1 - eff_draft_) * F_aero_)), and

c is equal to: 5.6986 * (1 + (6.13 * (1 - eff_draft_) * F_aero_)) - P_met og_

The discriminant, d, is equal to (b^2^ - 4ac). The quadratic equation can then be solved for v:

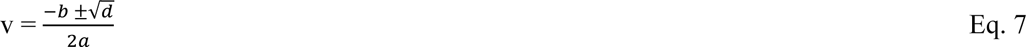

The next step is to solve Equation 5 for P_met og_ based on an actual race performance and the inferred drafting effectiveness. Then, it is possible to explore how much velocity would increase with greater drafting effectiveness values.

Using that approach, we focused on the 1 mile world record performance of Faith Kipyegon in Monaco on July 21, 2023 (https://www.youtube.com/watch?v=ndA9V4XBLvo). The ambient temperature was 29°C and the relative humidity was 57%. The barometric pressure at the start time (20:35 pm) was 751.8 mmHg (https://world-weather.info/forecast/monaco/monte_carlo/july-2023/) equating to an air density of 1.146 kg/m^3^ (https://www.calctool.org/atmospheric-thermodynamics/air-density). A trackside windsock visible in the broadcast video of the race indicated the wind was nearly dead calm. Kipyegon started on the inside of lane 1. Because 1 mile is 1609.344 m, on the Monaco 400 m per lap track, the starting line was 9.344 m before the finish line so that “Lap 1” was 409.344 m.

At the start, two designated pacemakers sprinted ahead in single file but there was a large spacing of ∼4 m between Kipyegon and the second pacemaker through ∼300 m. The spacing narrowed to about 3 m from ∼300 to ∼409 m. We utilized the 1.0 m spacing between the Wavelight pacing lights visible in the broadcast video to estimate the spacing between the pacer and Kipyegon. At ∼409 m, the spacing was reduced to ∼2.5 m and maintained to ∼809 m, with the first pacemaker exiting at ∼700 m. From ∼809 to ∼909 m, the spacing was ∼2 m and at 909 m, the remaining pacemaker moved out to lane 2, no longer providing any substantial drafting and then left the race at ∼1009 m. Kipyegon ran the remainder of the mile solo. According to the World Athletics website (https://monaco.diamondleague.com/programme-resultats/resultats-archives/), Kipyegon reached the 409.344 m mark in 1:02.60 and the 809.344 m mark in 2:04.60. Thus, her average velocity for Lap 1 (0 to 409.344 m) was 6.539 m/s and 6.452 m/s for Lap 2 (409.344 to 809.344 m). Kipyegon reached 1209.344 m in 3:06.80 and thus Lap 3 time was 62.20 sec which equates to a velocity 6.431 m/s. She then ran Lap 4 (1209.344 to 1609.344 m) in 60.84 sec or an average velocity of 6.575 m/s. Using those pacer-designated runner spacings and two previous studies of drafting effectiveness (Marro et al., 2022 and Schickhofer & Hanson, 2021), we estimated the impeding aerodynamic forces and hence Kipyegon’s total metabolic power during her record-breaking run using Equation 5. Although other previous studies have estimated drafting effectiveness for one specific pacer-designated runner spacing, only Marro et al. (2022) and Schickhofer & Hanson (2023) estimated drafting effectiveness across a range of spacings. We provide these calculations as a spreadsheet (Appendix 1) so that the reader can explore different assumptions. In these calculations, we calculate the mile times if Kipyegon was provided with optimal drafting for one, two, three and four laps. In the discussion section, we suggest options for how that drafting might be achieved.

Marro et al. (2023) used scaled manikins in a small, instrumented wind tunnel to estimate drag forces. Both Marro et al. (2023) and Schickhofer & Hanson (2022) denoted the configuration of one pacer in front of the designated runner as Formation 1 (Figure 1). Marro et al.’s data suggest that in Formation 1, with a pacer-designated runner spacing of ∼3-4 m, the drafting effectiveness was about 22% for Kipyegon’s Lap 1, and for Lap 2, the spacing of ∼2.5 m provided about a 30% drafting effectiveness. From 809-909 m, the spacing was ∼2 m and Marro et al.’s findings indicate a drafting effectiveness of about 32% but then as soon as the pacer moved to Lane 2 (initiated at 909 m), the drafting effectiveness likely went to zero. If we average 100 m at 32% and 300 m at 0%, we get 8% average drafting effectiveness for Lap 3. Since Kipyegon ran Lap 4 solo, the drafting effectiveness was 0. Marro et al. also estimated that when the pacer-designated runner spacing is 1.3 m, drafting effectiveness would be 39.5%. We used that 39.5% value in the spreadsheet to calculate how fast Kipyegon would be able to run with such optimal drafting for the full one mile race distance (see Appendix 1).

**Figure 1.**
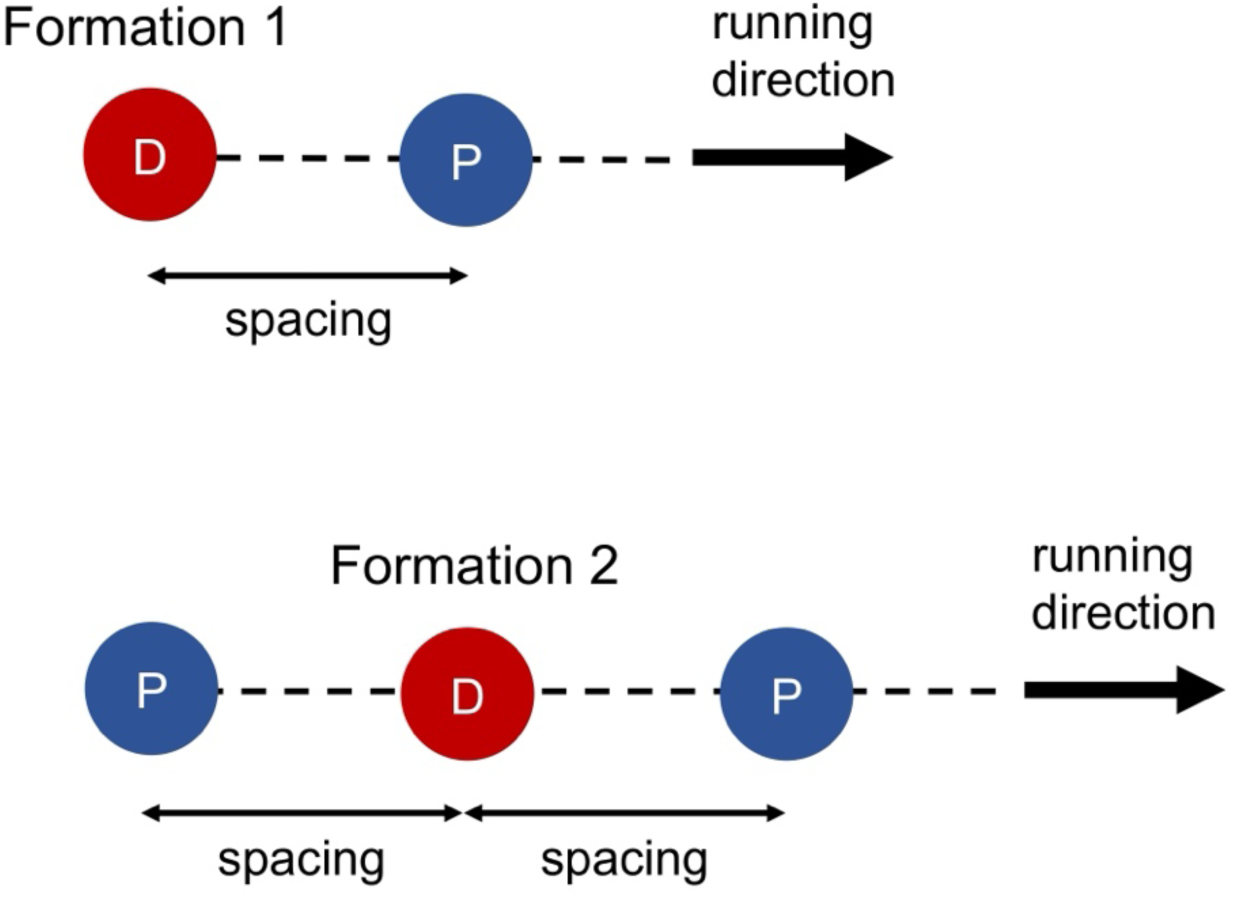
Formations 1 and 2. Red circles with the letter D represent the designated runner and blue circles with the letter P are pacers. Spacing is the distance between the designed runner and a pacer. Adapted from Schickhofer & Hanson (2021) and Marro et al. (2022).

**Figure 2.**
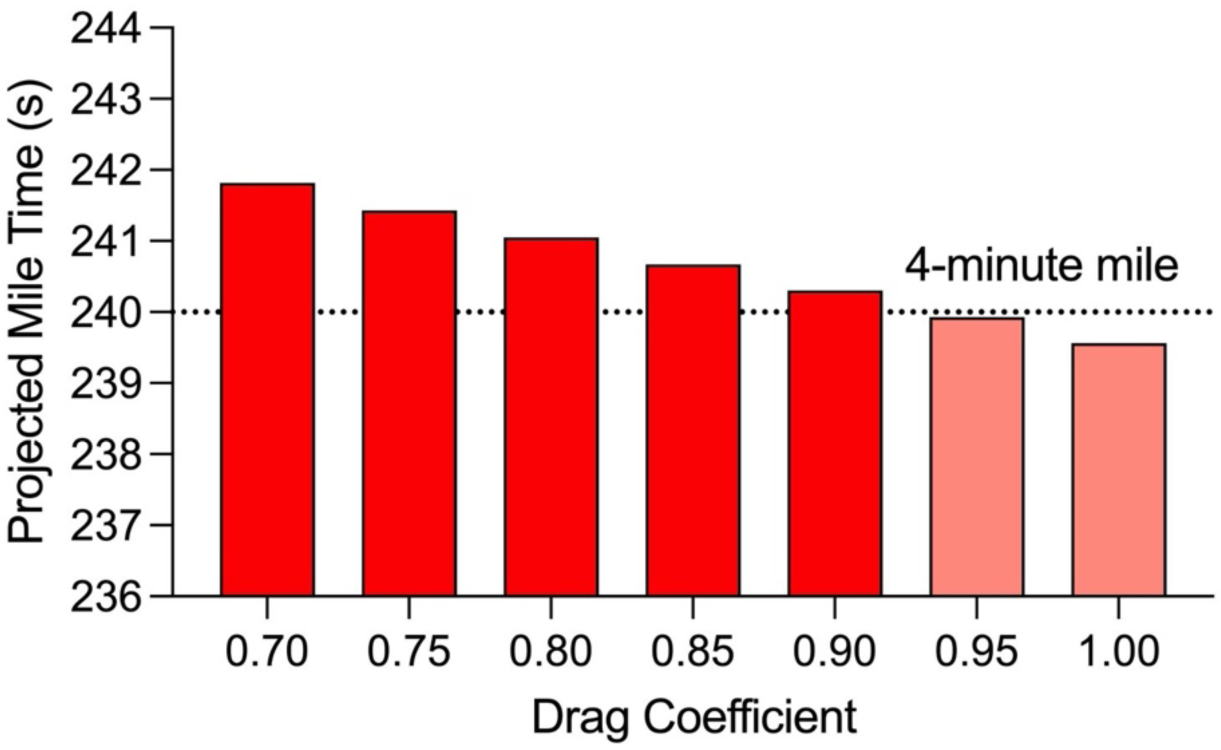
Predicted mile time changes were small in response to substantial changes in C_d_.

Using CFD, Schickhofer & Hanson (2021) found considerably greater drafting effectiveness values. Their simulation suggests that in Formation 1, with a spacing of ∼3-4 m, the drafting effectiveness would be ∼43% for Lap 1 and ∼48% for Lap 2. For 809 to 909 m, Schickhofer & Hanson predict ∼50% drafting effectiveness which when averaged with zero drafting for ∼909 to 1209 m equates to 12.5% drafting effectiveness for Lap 3. Again, Lap 4 had zero drafting. For Formation 1, with 1.2 m spacing, Schickhofer & Hanson found that drafting effectiveness is 70.1%. Again, we used that value in the spreadsheet to calculate how fast Kipyegon would be able to run with better drafting for the full one mile race distance. Further, Schickhofer & Hanson also simulated drafting effectiveness for one pacer 1.2 m in front combined with one pacer 1.2 m directly behind the designated runner (Formation 2) (Figure 1). That formation increases drafting effectiveness to 75.6% and we also made our calculations using that value.

We have retained a precision of 0.01 s (approximately 0.004%) in our calculations. We recognize that as being potentially misleading since our assumptions certainly have more than 1 s of uncertainty, which is ∼0.4%. Our justification follows here. We began our analysis with World Athletics times which are recorded to the 0.01 s. Rather than sequentially rounding the numbers in our calculations, we retained that precision. Running one mile (1609.344 m) in exactly four minutes (240.00 s) is an average velocity of 6.7056 m/s. Running a mile in 240.01 s equates to a velocity of 6.7053 m/s, and a 4:00.01 mile would not be as historically notable. That is, a difference of 0.01 second (or 0.001 m/s) is relevant in this situation. Thus, we have used 4 significant figures (e.g. 6.705 m/s) in all velocity calculations.

## 3. Results

We encourage the reader to utilize Table 2 and the spreadsheet provided in Appendix 1 through the following examples. For Lap 1 (409.344 m), Kipyegon’s actual time was 62.60 s or an average velocity of 6.539 m/s. Substituting that velocity into Eq. 5 and using a drafting effectiveness value of ∼22% for a pacer-designated runner spacing of ∼3-4 m from Marro et al. (2022) gives a metabolic power of 35.01 W/kg. If we keep that metabolic power value but improve the drafting effectiveness to 39.5% as Marro et al. (2022) suggest for a spacing distance of 1.3 m in Formation 1, then we can solve Equation 5 for the faster v. In this case, Lap 1 velocity increases to 6.618 m/s or a time for Lap 1 of 61.86 s, an improvement of 0.74 s. For Lap 2 (400 m), the Monaco time was 62.00 s (6.452 m/s) and drafting effectiveness was ∼30% which equates to a metabolic power of 33.86 W/kg. Increasing drafting effectiveness to 39.5% would allow velocity to increase to 6.493 m/s and a Lap 2 time of 61.60 s, an improvement of 0.40 s. Thus, providing improved drafting for just two laps yields a mile time of 4:06.82. For Lap 3 (400 m), the Monaco time was 62.20 s (6.431 m/s) and drafting effectiveness averaged ∼8% yielding a metabolic power of 34.52 W/kg. Increasing drafting effectiveness to 39.5% would allow velocity to increase to 6.566 m/s and a Lap 3 time of 60.92 s, an improvement of 1.28 s. Thus, if improved drafting was provided for three laps, Kipyegon’s mile time would be 4:05.22. For Lap 4 (400 m) in Monaco, the time was 60.84 s (6.575 m/s) and there was no drafting. Thus, according to Equation 3, the metabolic power was 36.27 W/kg. Increasing drafting effectiveness to 39.5% would allow velocity to increase to 6.753 m/s and a Lap 4 time of 59.23 s, an improvement of 1.61 s. Summing those four lap times equals 4:03.61 for the full mile.

**Table 1.** Drafting effectiveness values based our estimates of the spacing between the pacers and Kipyegon for each lap of her mile world record race in 2023 in Monaco. The corresponding drafting effectiveness values were read from the graphs in Marro et al. (2023) and Schickhofer & Hanson (2022). Improved drafting effectiveness values were taken from those same studies for one pacer in front with 1.3 and 1.2 m spacings respectively. We assumed that pacing could provide drafting for the full mile distance.

| Laps | Monaco Spacing | Monaco Drafting Effectiveness<sup>1</sup> | Improved Drafting Effectiveness<sup>1</sup> | Monaco Drafting Effectiveness<sup>2</sup> | Improved Drafting Effectiveness<sup>2</sup> |
| --- | --- | --- | --- | --- | --- |
| Lap 1 | ~3-4 m | 22% | 39.5% | 43% | 70.1% |
| Lap 2 | ~2.5 m | 30% | 39.5% | 48% | 70.1% |
| Lap 3 | ~2.0 m, N/A | 8% | 39.5% | 12.5% | 70.1% |
| Lap 4 | N/A | 0% | 39.5% | 0% | 70.1% |

N/A (not applicable) indicates that there was zero drafting from ~909 m through the finish. 1 - Marro et al., (2023); 2 - Schickhofer & Hanson (2022)

**Table 2.** Lap times (min:sec) calculated for various drafting assumptions. We used: a metabolic cost factor for horizontal resistive force of 6.13% per body weight (da Silva et al., 2022), air density of 1.146 kg/m^3^, coefficient drag of 0.90 and drafting effectiveness values from Marro et al. (2023) for the pacer to designated runner spacings observed on the broadcast video of Kipyegon’s world record race in Monaco. These calculations all involve Formation 1.

| Scenario (drafting effectiveness %) | Lap 1 | Lap 2 | Lap 3 | Lap 4 | Final |
| --- | --- | --- | --- | --- | --- |
| Monaco (4 Laps: 22, 30, 8, 0%) | 1:02.60 | 1:02.00 | 1:02.20 | 1:00.84 | 4:07.64 |
| 2 Laps drafting (39.5, 39.5, 0, 0%) | 1:01.86 | 1:01.60 | 1:02.52 | 1:00.84 | 4:06.82 |
| 3 Laps drafting (39.5, 39.5, 39.5, 0%) | 1:01.86 | 1:01.60 | 1:00.92 | 1:00.84 | 4:05.22 |
| 4 Laps drafting (39.5, 39.5, 39.5, 39.5%) | 1:01.86 | 1:01.60 | 1:00.92 | 59.23 | **4:03.61** |
| No Drafting (all 4 Laps: 0%) | 1:03.53 | 1:03.24 | 1:02.52 | 1:00.84 | 4:10.14 |
| No Air resistance (all 4 Laps: 100%) | 59.26 | 59.07 | 58.44 | 56.74 | 3:53.51 |

If instead, we use the drafting effectiveness values from Schickhofer & Hanson (2021), we calculate that Kipyegon could run substantially faster. For Lap 1, the Monaco velocity was 6.539 m/s and using Schickhofer & Hanson’s Formation 1 drafting effectiveness value of 43% gives a metabolic power of 34.15 W/kg. Improving the drafting effectiveness to 70.1% as Schickhofer & Hanson suggest for Formation 1 with a pacer-designated runner spacing of 1.2 m, Lap 1 velocity increases to 6.665 m/s or a time of 61.42 s, an improvement of 1.18 s. For Lap 2 (400 m), Kipyegon’s Monaco velocity was 6.452 m/s and with a drafting effectiveness value of 48% we calculate a metabolic power of 33.16 W/kg. Improving drafting effectiveness to 70.1% increases Lap 2 velocity to 6.551 m/s or a time of 61.06 s, an improvement of 0.94 s. Thus, providing drafting for two laps yields a time of 4:06.03. For Lap 3 (400 m), Kipyegon’s Monaco velocity was 6.431 m/s which combined with a drafting effectiveness value of 12.5% gives a metabolic power of 34.34 W/kg. Improving drafting effectiveness to 70.1% increases Lap 3 velocity to new 6.684 m/s or a time of 59.84, an improvement of 2.36 s. Thus, providing drafting for three laps yields a mile time of 4:03.16. For Lap 4 (400 m), Kipyegon’s Monaco velocity was 6.575 m/s and with zero drafting that required a metabolic power of 36.27 W/kg. Improving drafting effectiveness to 70.1% increases Lap 4 velocity to 6.899 m/s or a time of 57.98 s, an improvement of 2.86 s. Thus, for the full mile distance, her time would be 4:00.30. Keeping all the same assumptions but changing drafting effectiveness to 75.6% for four laps of drafting in Formation 2 yields a time of 3:59.37, breaking the 4-minute mile barrier!

Next, we used the spreadsheet to calculate how fast Kipyegon could have run in two hypothetical situations: solo with no drafting and with zero air resistance, such as on a treadmill. If we retain the Marro et al. (2022) Formation 1 drafting effectiveness values for Monaco, but zero out the hypothetical drafting effectiveness values, the calculations predict that running solo, Kipyegon could run 4:10.14 vs. her actual 4:07.64, a difference of 2.50 s. Retaining the Schickhofer & Hanson Formation 1 drafting effectiveness values for Monaco and again zeroing out the hypothetical drafting effectiveness, we calculate that she would have run 4:12.04 solo, a difference of 4.40 s. On the other hand, if we retain the Marro et al. drafting effectiveness values for Kipyegon in Monaco but input 100% hypothetical drafting effectiveness in Formation 1, we calculate a mile time of 3:53.51. Using Schickhofer & Hanson Formation 1 values, we obtain 3:55.24. In both examples, the discrepancy between the two predictions is due to Marro et al.’s finding of generally lower drafting effectiveness values than Schickhofer & Hanson.

## 4. Discussion

Our calculations support the hypothesis that with optimal aerodynamic drafting, an elite female athlete could break the 4-minute mile barrier. Combining the Schickhofer & Hanson (2021) aerodynamic data with our physiological model, we calculate that a female athlete would need to be provided with at least 71.9% drafting effectiveness for the full mile distance to run 3:59.99 (See Table 3). Schickhofer & Hanson found that a drafting effectiveness of 75.6% could be achieved in Formation 2, i.e. one pacer 1.2 m in front of a designated athlete combined with a second pacer 1.2 m behind the designated athlete. With 75.6% drafting effectiveness, our calculations predict that Kipyegon could run 3:59.37. Coincidentally, that is essentially the same time that Bannister ran in his first 4-minute mile. Using our same approach but with the scaled wind tunnel estimates of Marro et al., (2022) and four laps of drafting, we calculated a best mile time of 4:03.61. It is not yet clear whether Schickhofer & Hanson or Marro et al.’s aerodynamic estimates are most accurate.

**Table 3.** Lap times (min:sec) calculated for various drafting assumptions. We used: a metabolic cost factor for horizontal resistive force of 6.13% per body weight (da Silva et al., 2022), air density of 1.146 kg/m^3^, drag coefficient of 0.90 and drafting effectiveness values from Schickhofer & Hanson (2022) for the pacer to designated runner spacing observed on broadcast video of Kipyegon’s world record race in Monaco. The 70.1% drafting effectiveness is for Formation 1 and the 75.6% drafting effectiveness is for Formation 2. We used the spreadsheet to iteratively solve for the 71.9% drafting effectiveness which is the minimum that would allow a sub-4-minute mile.

| Scenario (drafting effectiveness %) | Lap 1 | Lap 2 | Lap 3 | Lap 4 | Final |
| --- | --- | --- | --- | --- | --- |
| Monaco (4 Laps: 43, 48, 12.5, 0%) | 1:02.60 | 1:02.00 | 1:02.20 | 1:00.84 | 4:07.64 |
| 2 Laps drafting (70.1, 70.1, 0, 0%) | 1:01.42 | 1:01.06 | 1:02.71 | 1:00.84 | 4:06.03 |
| 3 Laps drafting (70.1, 70.1, 70.1, 0%) | 1:01.42 | 1:01.06 | 59:84 | 1:00.84 | 4:03.16 |
| 4 Laps drafting (70.1, 70.1, 70.1, 70.1%) | 1:01.42 | 1:01.06 | 59:84 | 57:98 | **4:00.30** |
| 4 Laps drafting (75.6, 75.6, 75.6, 75.6%) | 1:01.18 | 1:00.83 | 59:61 | 57:75 | **3:59.37** |
| No Drafting (all 4 Laps: 0%) | 1:04.46 | 1:04.03 | 1:02.71 | 1:00.84 | 4:12.04 |
| 4 Laps drafting (71.9, 71.9, 71.9, 71.9%) | 1:01.34 | 1:00.93 | 59.77 | 57:90 | **3:59.99** |
| No Air resistance (all 4 Laps: 100%) | 1:00.11 | 59.78 | 58.61 | 56.74 | 3:55.24 |

Since our calculations depend on several assumptions, we performed sensitivity tests using the Schickhofer & Hanson (2021) drafting effectiveness values. We assumed a drag coefficient of 0.90 as per (Hill, 1927; Davies, 1980; and Marro et al. 2023) but C_d_ values in the literature range from 0.80 to 1.1. We found that in Formation 1, decreasing C_d_ to 0.80 would add less than a second to the final time and increasing C_d_ to 1.00 would reduce the final time by less than a second (see Table 4). Such changes in C_d_ are large, yet the model predictions were robust.

**Table 4.** The effects of varying drag coefficient on running performance over each of the four laps. In the literature, drag coefficient values range from 0.8 to 1.1 for human solo running (Davies, 1980; Pugh, 1971; Hill, 1927; Shanebrook & Jaszczak, 1976). Our calculations used a metabolic cost factor for horizontal resistive force of 6.13% per body weight (da Silva et al., 2022), 70.1% drafting effectiveness from Schickhofer & Hanson (2022) and air density of 1.146 kg/m^3^.

| Drag coefficient | Lap 1 | Lap 2 | Lap 3 | Lap 4 | Final |
| --- | --- | --- | --- | --- | --- |
| 0.70 | 1:01.67 | 1:01.26 | 1:00.33 | 58:56 | 4:01.82 |
| 0.75 | 1:01.61 | 1:01.21 | 1:00.21 | 58:41 | 4:01.43 |
| 0.80 | 1:01.54 | 1:01.16 | 1:00.08 | 58:26 | 4:01.05 |
| 0.85 | 1:01.48 | 1:01.11 | 59:96 | 58:12 | 4:00.67 |
| 0.90 | 1:01.42 | 1:01.06 | 59:84 | 57:98 | 4:00.30 |
| 0.95 | 1:01.36 | 1:01.01 | 59:72 | 57:83 | 3:59.93 |
| 1.00 | 1:01.30 | 1:00.96 | 59:60 | 57:69 | 3:59.56 |

As per da Silva et al. (2022), we assumed a value of 6.13% change in metabolic power per BW of horizontal resistive force. That value was the mean empirical finding but individual values ranging from 4.17 to 8.14%. We do not have any empirical measurements for Kipyegon and she may be an average, high or low responder to horizontal resistive forces. Thus, we tested the sensitivity of our calculations by assuming a variety of metabolic cost factors from 4 to 8% (see Table 5 and Figure 3). We found that decreasing the metabolic cost factor from 6.13%/BW to 4.0%/BW increased the mile time by ∼2.5 s. Conversely, we calculated that increasing the metabolic cost factor to 8%/BW, would decrease the mile time by ∼2 s. We recognize that the fastest running speed we studied in da Silva et al., (2022), 4.4 m/s, is slower than 4-minute mile pace (∼6.5 m/s). But, given that the metabolic cost factor did not substantially differ from 3.3 to 4.4 m/s, we have some confidence in applying the average da Silva et al. metabolic factor (6.13%/BW) to 6.5 m/s running, even though it is an extrapolation.

**Figure 3.**
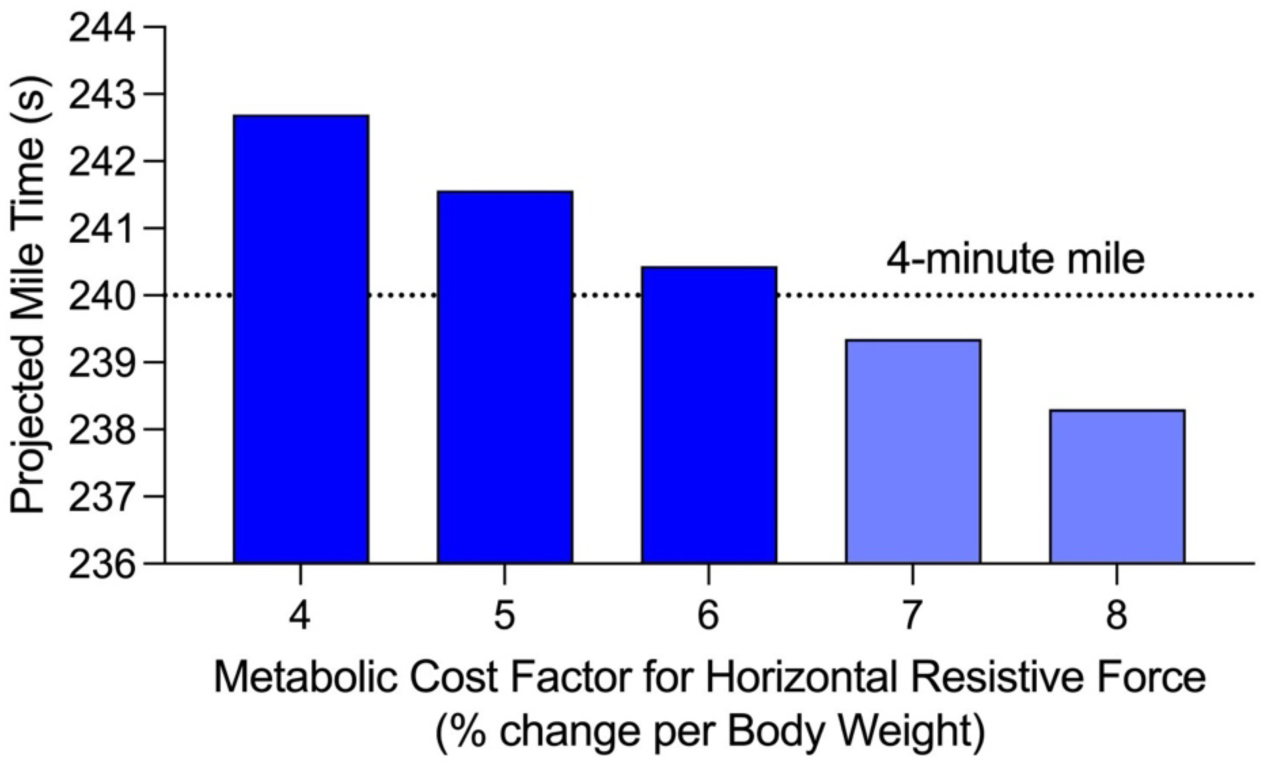
Predicted mile time changes were relatively small in response to substantial changes in the metabolic cost factor.

**Table 5.** The effect of varying the metabolic cost factor for horizontal resistive force on running performance over the four laps. Calculations used 70.1% drafting effectiveness, a drag coefficient drag of 0.90 and air density of 1.146 kg/m^3^ for all four laps.

| Increase in metabolic cost (%) | Lap 1 | Lap 2 | Lap 3 | Lap 4 | Final |
| --- | --- | --- | --- | --- | --- |
| 4 | 1:01.81 | 1:01.38 | 1:00.61 | 58:90 | 4:02.70 |
| 5 | 1:01.63 | 1:01.23 | 1:00.24 | 58:46 | 4:01.56 |
| 6 | 1:01.44 | 1:01.08 | 59:89 | 58:03 | 4:00.44 |
| 7 | 1:01.26 | 1:00.94 | 59:54 | 57:62 | 3:59.35 |
| 8 | 1:01.09 | 1:00.80 | 59:20 | 57:21 | 3:58.30 |

Our frontal area estimate could also have affected our calculations. We used surface and frontal area equations that were developed for males (Du Bois & Du Bois, 1916; Pugh, 1970) but of course, females and males are not all the same shape. Fortunately, Schickhofer & Hanson (2022) provided empirical data that allow us to test our frontal area estimate for Kipyegon. They measured the height and body mass of 59 female athletes and measured their frontal area from photos. They used those data to calculate the frontal area of a 55 kg, 1.65 m tall female athlete as 0.44 m^2^. If we input 55 kg and 1.65 m into the Du Bois & Du Bois and Pugh equations, we calculate a remarkably similar projected frontal area (0.43 m^2^). Such close correspondence gives us confidence that our frontal area estimate for Kipyegon is reasonable and accurate.

When Kipyegon set the world record for the mile in 2023 in Monaco, the air had a low density due to the high temperature and humidity. Hotter or more humid conditions would provide a lower air density which might be advantageous, but the thermoregulation challenges would likely make it more difficult to run a 4-minute mile (Yamashita et al., 2024). Similarly, running at altitude would reduce air resistance but the lower partial pressure of oxygen would have a much greater negative effect on aerobic capacity (Peronnet et al., 1991).

In practice, adequate aerodynamic drafting for a female 4-minute mile attempt could be achieved in several ways. First, the designated elite female could run a time trial with two elite males (i.e. established 4-minute milers) as designated pacers for the full distance in Formation 2. That would not conform to World Athletics eligibility regulations for a female-only world record. Second, a designated elite female could have elite female middle-distance runners as pacers in Formation 2 for the first two laps. Then, towards the end of the second lap, the first pair of pacers could move off the track and a second pair could pace the designated runner for the third and fourth laps. That would mimic the method pioneered in the Breaking2 project during which Eliud Kipchoge nearly broke the 2-hour marathon barrier. But again, World Athletics rules would not recognize such a performance as eligible for a world record because pacers are not allowed to enter a race enroute.

In all these scenarios, pacing lights on the track curbing would be helpful and such light systems (i.e. Wavelight Technologies, Nijmegen, The Netherlands) are now commonplace in elite competitions. Further, in any of these scenarios, the athletes would need to devote significant practice time to coordinate the choreography of their movements. The athletes would also need to practice minimizing the designated athlete-pacer spacing. Any attempt to break the 4-minute mile barrier should target a location/day/time with minimal wind. Both Marro et al. and Schickhofer & Hanson assume that the designated athlete and the pacers have the same body dimensions. However, Kipyegon is small in stature (1.57 m) compared to many elite female 800 m specialists (World Athletics, 2024). For example, the Tokyo Olympic champion, Athing Mu has a height of 1.78 m and the 2024 Paris Olympic champion, Keely Hodgkinson is 1.70 m tall. If tall pacers were selected, Kipyegon might experience better drafting effectiveness than we have assumed, though a taller pacer with longer legs might make it more difficult to maintain close spacing. Finally, more effective, pacer formations are also possible. The arrowhead or reverse arrowhead pacer formations used to break the 2-hour marathon could be implemented on the unusual track configuration of Franklin Field at the University of Pennsylvania (https://en.wikipedia.org/wiki/Franklin_Field). On that track, the designated runner could run in lane 4, which is 400 m in circumference, with pacers running in an arrowhead formation in lanes 1-3 and 4-7.

A possible limitation to our method/model is that several studies we rely on, did not include any female participants. Specifically, Batliner et al., (2018) only studied males. However, running economy (expressed per kg body mass) does not differ substantially between males and females (Daniels & Daniels, 1992). Further, even if there was a difference, our model only uses relative changes in velocity at a given metabolic power and it is not dependent on absolute values for metabolic power. We also rely on da Silva et al., (2022) who only studied male participants. However, the key factor from da Silva et al., is scaled to body weight and it is difficult to imagine why such a scaled factor would differ between e.g. a 60 kg male and a 60 kg female. Unfortunately, we are not alone; the field of exercise physiology needs to improve in terms of including female study participants (Cowley et al., 2021). An organized scientific effort to facilitate the first female 4-minute mile would be an excellent opportunity to study elite female runners and create excitement for more female-based studies in the field of exercise physiology.

## 5. Conclusions

Our calculations suggest that with greatly improved (but reasonable) aerodynamic drafting, the current record holder, Faith Kipyegon could break the 4-minute mile barrier. We find that she could feasibly run ∼3:59.37 with two teams of female pacers (one in front and one in the back) who change out at 800 m. Hopefully, Ms. Kipyegon can test our prediction on the track.

## Supporting information

Female 4 min mile spreadsheet

## Footnotes

* We calculated the body surface area (SA) for Roger Bannister and Faith Kipyegon with the Du Bois & Du Bois (1916) equation, where the height (H) is in cm and body mass (M) in kg: SA = 0.007184 H^0.725^ M^0.425^ Then, we calculated frontal area (A_f_) based on Pugh (1970): A_f_ = 26.6% SA.

